# Biological implications of a detailed repeat annotation in *Octopus vulgaris*

**DOI:** 10.64898/2026.03.03.709284

**Authors:** Maegwin Bonar, Tyler A. Elliott, Mirza A. M. Ahmadi, Karl Cottenie, Stefan Linquist

## Abstract

Octopuses are phenotypically distinctive organisms, and recent genomic work raises questions about the contributions of transposable elements (TE) to their genomic architecture. We leveraged a robust repeat annotation pipeline, in combination with manual and automated curatorial techniques, to produce a more comprehensive repeat annotation of *Octopus vulgaris*. This revealed that ∼66% of the genome are repeats, in contrast to previous estimates of 43-50%. Whereas previous studies of TE expansion in *Octopus bimaculoides* identified two bursts of activity, 25 and 56 MYA, our re-annotation revealed four such expansions at 18, 25, 33, and 56 MYA. We further identified a landscape of TE hot- and cold spots. This much refined TE timescape and landscape will serve as a useful basis for understanding TE contributions to *O. vulgaris* evolution, and also for identifying factors contributing to variation in the TE community across genomic space and evolutionary time.

## Introduction

Octopuses are highly derived molluscs exhibiting a variety of novel traits not found outside the coleoid cephalopod lineage. Some of these phenotypic novelties appear to be associated with large-scale genome reshuffling events, gene duplications, and recent expansion of transposable elements (TEs) (Albertin et al., 2015; 2022; Yoshida et al., 2025). Transposable elements are mobile strands of DNA often containing repetitive sequences potentially involved in genome restructuring (Lönnig and Saedler 2002). Recent octopus genome annotations identify two periods of significant TE expansion, 56 and 25 MYA (Albertin et al 2015). Some TE insertions are associated with genes that underwent expansion in the octopus genome, and full-length LINE transcripts are expressed in octopus neurons (Petrosino et al., 2022). Taken together, these findings have been interpreted as evidence for TE involvement in generating phenotypic innovation in cephalopods (Albertin 2015, 2022) including complexity of their nervous systems; although there are alternative explanations for these patterns (Linquist et al. 2026).

Recently, an improved genome assembly of *Octopus vulgaris* has become available (Destanovic et al., 2023), providing opportunities for better annotation including the potential for understanding TE contributions to phenotypic evolution. While detailed curation is known to improve repeat annotation, including the detection of active TEs (Platt et al., 2016; Goubert et al., 2022), previous annotations of octopus genomes did not include curatorial work. In this study, we used a long-read chromosome-scale genome assembly for *O. vulgaris* and a robust annotation pipeline to generate a comprehensive repeat annotation for this species. We then explored the activity timescape and the distribution and abundance landscape of TEs. This revealed a greater diversity of TEs, more recent periods of expansion than previously documented in coleoids, and regions of hotspot/coldspot TE accumulation.

## Results & Discussion

### Repeat annotation summary

Repeat detection and annotation with Earl Grey and a custom pipeline of additional tools along with a curatorial process (see Materials and Methods) resulted in 66.25% of the *O. vulgaris* assembly annotated as repeats (Supplementary Table 1). The most abundant TE orders are LINE (12.9% of the genome) and SINE elements (11.6%) (Figure 1A). This concurs with a previous raw-data Genomescope estimate of 68.68% for *O. vulgaris* (Destanovic et al., 2023), but is considerably larger than similar, assembled estimates for *O. bimaculoides (*43-47% repeats) (Albertin et al., 2015) and in *O. sinensis* (42.3%) (Li et al., 2020). The greater percentage of repeats and diversity of TEs identified here, compared to previous studies that did not use curation or supplementary tools, demonstrate the advantages of these additional steps (see also Platt et al., 2016; Goubert et al., 2022).

**Figure 1.**
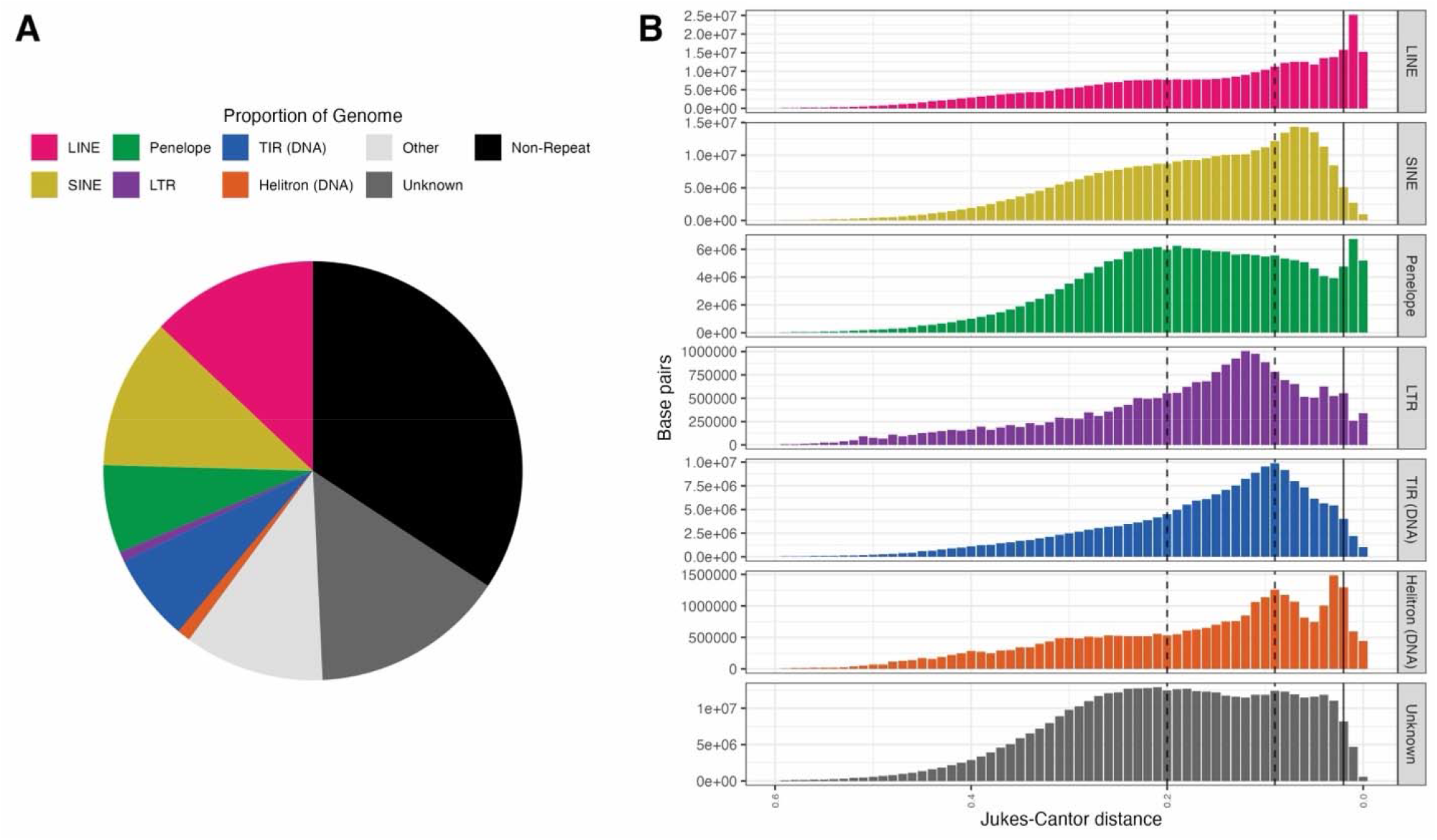
Repeat detection and insertion history of transposable elements (TEs) in *O. vulgaris* genome. A) Pie chart illustrating the proportion of base pairs in the *O. vulgaris* genome comprising the six TE orders (LINE, SINE, Penelope-like elements, LTR, TIR, Helitron), other repeats (i.e., simple repeats, microsatellites, low-complexity elements), unknown repeats, and non-repeat elements. B) Age of TEs measured in Jukes-Cantor distance. Solid black line represents the threshold for active elements, dashed black lines illustrate the time points in which Albertin et al. (2015) identified two major expansion waves for TEs in *O. bimaculoides* at Jukes-Cantor distance 0.09 and 0.2 respectively.

### TE timescape

Previous studies in *O. bimaculoides* (Albertin et al. 2015) identified two periods of TE expansion estimated at 25 mya (Jukes-Cantor (JC) distance 0.09) and 56 mya (JC distance 0.2). We dated all TE insertions using JC distances between individual copies and their consensus sequence (Jukes and Cantor, 1969; Lander et al., 2001). This revealed four periods of TE expansion, two of which correspond with previous work by Albertin et al. (2015). These timepoints correspond to expansions of TIRs and Helitrons at 25 million years (JC distance 0.09), and with Penelope-like elements and LINEs expanded at 56 million years (JC distance 0.2) (Figure 1B). The two additional expansions identified in our study involved SINEs and LINEs at 18 million years (JC distance 0.065) and an expansion of LTR elements at 33 million years (JC distance 0.12). *O. vulgaris* is estimated to have diverged from *O. bimaculoides* between 13.3 and 33.5 mya (Huang et al., 2022). These results suggest that TEs have been more active in the *O. vulgaris* genome than previously reported, although some of discrepancy could be due to species differences.

We defined recently active TEs as insertions ≤ 2% divergent from consensus sequences and ≥ 90% the length of the consensus sequence. Over 11000 insertions from 1298 families were identified as recently active, which included members from all TE orders and unknown repeats (Table 1). The recently active insertions comprised approximately 19 Mb, or 0.0068 % of genome and 0.01% of TEs. The diversity of TE orders in the recently active insertions contrasts with the human genome which, although similar in size, only retains a limited number of active TEs from LINEs and SINEs (Mills et al., 2007). These findings reveal a large diversity of active elements in *O. vulgaris*.

**Table 1.**
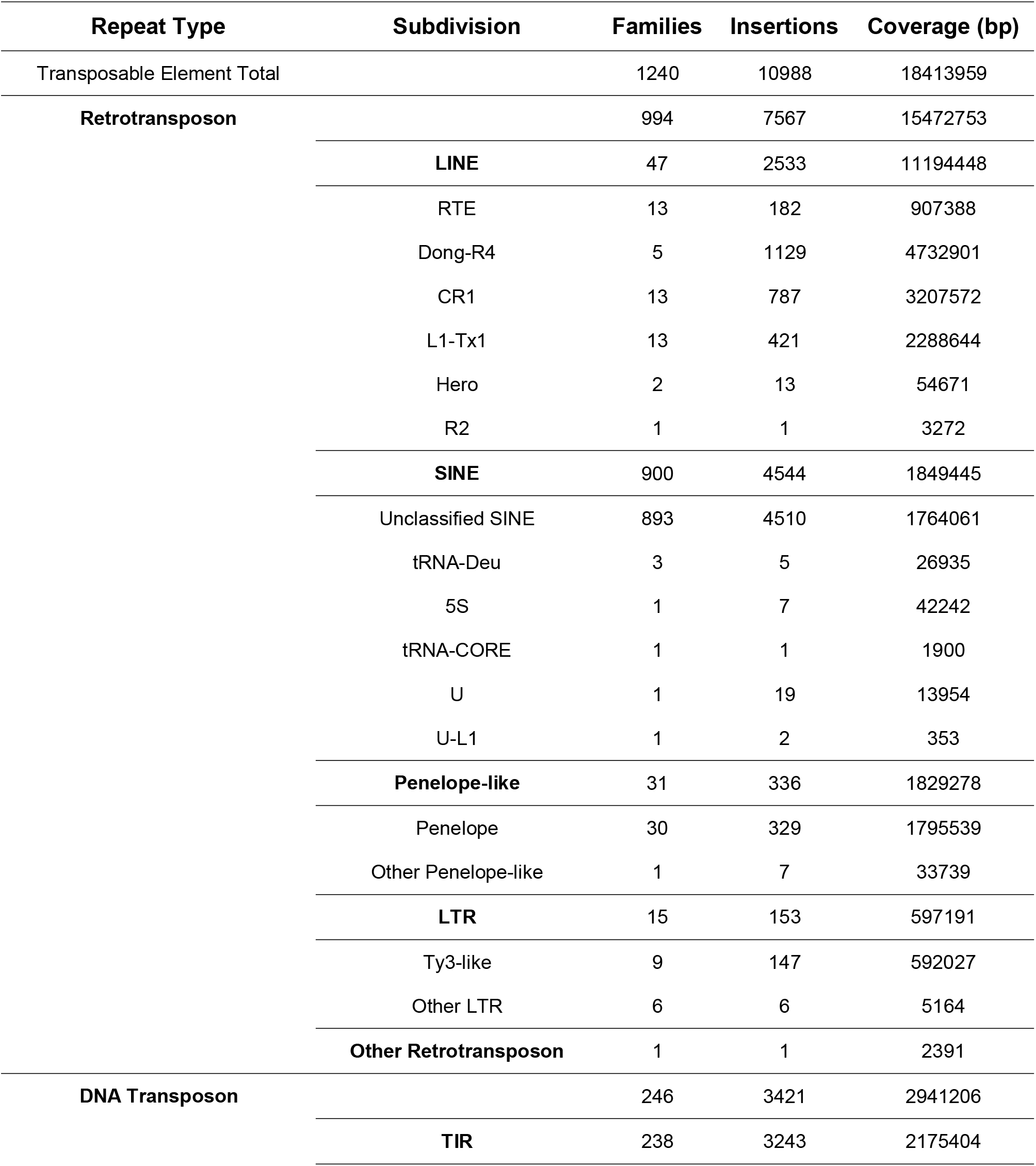
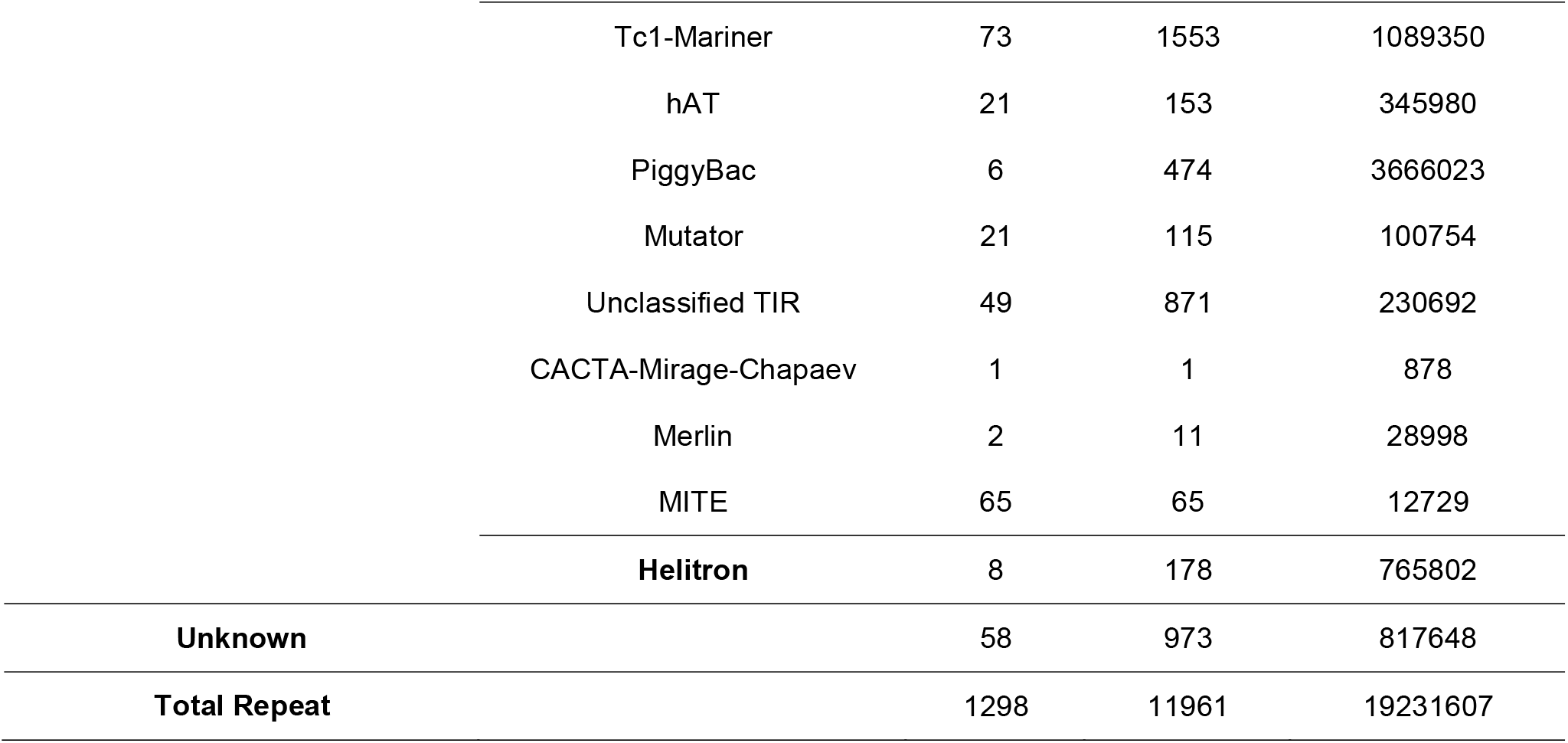
Recently active insertions in the *Octopus vulgaris* genome. Insertions were defined as recently active if elements had 2% or less divergence from consensus sequence and 90% or greater length of the consensus sequence.

RTE and Dong-R4 LINEs have been previously identified as possibly being transpositionally active in the brain of *O. vulgaris* and *O. bimaculoides* (Petrosino et al., 2022). Petrosino et al. (2022) did not report sequences of the active elements in the brain, and thus we were unable to determine if these were part of our recently active RTE and Dong-R4 LINE families. In contrast, we identified a combined total of 18 potentially active families and over 1200 potentially active individual insertions of RTE and Dong-R4 LINEs (Table 1). If TEs play some organism-functional role in the octopus brain, then it is a much larger diversity than previously reported.

### TE landscape

TEs were unevenly distributed across the *O. vulgaris* genome, as indicated by a high standard deviation in TE base pairs per 100kb genomic window (mean = 53.83kb/100kb, SD = 8.65kb/100kb). We identified 288 hotspots of TE accumulation in the *O. vulgaris* genome (Figure 2), defined as 100kb windows with a TE content above a 99% area under the curve (AUC) cut-off (see Methods; Baril and Hayward 2022). These hotspots contained a TE density of at least 1.4 times higher than average (mean = 53.83kb, lowest hotspot density = 76.02kb, highest hotspot density = 93.83kb). Recently active elements occurred in 61.8% of the hotspot (178 windows; density of active elements range 0.1kb - 36.8kb). We identified 278 TE coldspots across the genome, defined as 100kb windows with TE content below a 1% AUC cut-off, where TE density was at least 1.6 times lower than average (lowest coldspot density = 1.37kb, highest coldspot density = 32.73kb). Active elements were present in low densities in 17.3% of coldspots (48 windows; density of active elements range 0.1kb - 0.6kb). Other studies have identified the genome-level equivalent of ecological “niches” in maize (Stitzer et al., 2021), and these hotspots could be an equivalent phenomenon in the octopus genome.

**Figure 2.**
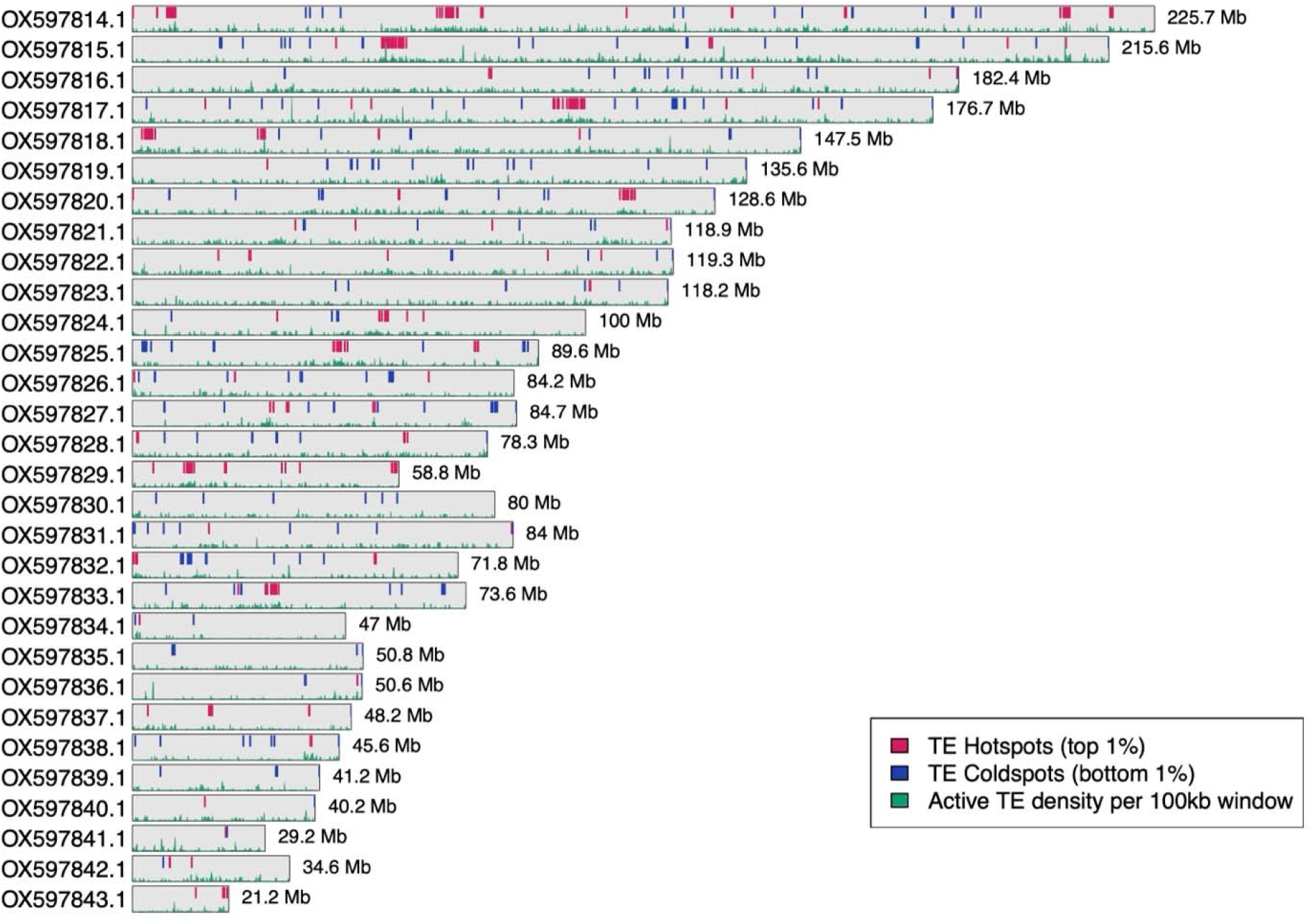
Karyoplot illustrating transposable element (TE) landscape for the *O. vulgaris* genome. Red and blue boxes indicate the position of 100kb windows with TE content above a 99% area under the curve (AUC) cut-off and (hotspots) and windows with TE content below the bottom 1% AUC cut-off (coldspots). Green bars illustrate the location of active TE elements, each bar representing 100kb windows and the height of bars represents the density of active elements in that window.

## Conclusion

We provide a detailed annotation of TEs in the *O. vulgaris* genome and an analysis of the diversity and distribution of TEs including recently active elements. This reveals a greater abundance and diversity of TEs than previously estimated, and multiple historical TE expansions. Our study lays the foundation for future work with *O. vulgaris* investigating the mechanisms driving TE expansion and distribution and the role TEs play in the evolution of this host species.

## Materials and Methods

### Data

We analyzed the xcOctVulg1.2 (GCA_951406725.2) *O. vulgaris* genome assembly, encompassing 2.8 Gb assembled from Oxford Nanopore and 10x Chromium reads, with a gene and lncRNA annotation (Destanovic et al., 2023). This was assembled into 30 chromosome-scale scaffolds, in concordance with karyotype information known for this species (Gao and Natsukari, 1990). Reference mollusc TEs were gathered from various sources, including Repbase version 23.02 (Bao et al., 2015), Dfam release 3.7 (Storer et al., 2021) and others detailed in Supplementary Material.

### Repeat detection

We used Earl Grey v5.0.0 (Baril et al., 2024) for repeat detection using the largest scaffolds (∼ 1Gb) in accordance with the recommendations from Jamilloux et al. (2017) for a larger genome that is less computationally taxing. Included in Earl Grey is RepeatModeler2 v2.0.5 (Flynn et al., 2020) for *de novo* repeat detection and classification, LTR_FINDER version v1.0.7 (Xu and Wang, 2007; Ou and Jiang, 2019) for LTR element detection, and a BLAST-extract-align-trim process for simple curation of repeats. We applied additional tools, derived from the TE-Atlas pipeline (https://github.com/mirzaahmadi/TE-Atlas) including HELIANO v1.0.2 (Li et al., 2024), AnnoSINE v2 (Li et al., 2021), HiTE v3.3.2 (Hu et al., 2024), and MiteFinderII (Hu et al., 2018). We merged the resulting TE libraries from all tools and clustered using CD-HIT-EST v4.8.1 (Li and Godzik, 2006) at 80% sequence identity to group elements into TE families (Wicker et al., 2007).

### Repeat curation

We interrogated the classification of sequences in the TE library using TEsorter version 1.4.6 (Zhang et al., 2022), and TE-Aid (Goubert et al., 2022). We detected and removed host gene contamination using Pfam_scan (Finn et al., 2014) and Pfam database release 35 (Mistry et al., 2021). We curated consensus sequences for TE families with TEtrimmer version 1.5.4 (Qian et al., 2025) to collect hits from the assembly for each consensus, align hits, extend or trim the ends of consensus sequences based upon the alignment, orient consensus sequences based on the direction of coding regions, and remove redundant consensus sequences.

### Repeat annotation

We input the clustered, combined and curated library of reference and *de novo* repeats into the Earl Grey AnnotationOnly command to annotate repeats in the complete *O. vulgari*s assembly (2.8 Gb). Earl Grey also detects tandem repeats with TRF version 4.09.1 (Benson, 1999), and repeats from the total library and TRF were annotated in the assembly using RepeatMasker version 4.1.5. Close or overlapping repeats of the same consensus sequence were merged using RepeatCraft version 1.0 (Wong and Simakov, 2019) to produce a final repeat annotation. TE annotations shorter than 100 bp were filtered out to exclude potential spurious annotations for summary statistics (Baril and Hayward, 2022; Baril et al., 2024).

### TE timescape

We defined recently active TEs as elements with ≤ 2% divergence from consensus sequence and ≥ 90% length of the consensus sequence. Divergence from consensus was estimated using the Kimura 2-parameter (K2P) mutation model (Kimura, 1980) which is part of the standard Early Grey output. We then determined the age of TE insertions using the K2P divergence from consensus and adjusted the distances for multiple substitution using the Jukes-Cantor (JC) formula:

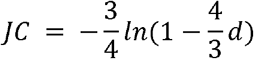

Where d is the divergence distance estimated by Earl Grey. We converted the JC distances to time in millions of years using the estimate for synonymous neutral substitutions per million years for *O. bimaculoides* (dS = 0.0036; Albertin et al. 2015) given the recent divergence between the two species and that there is no estimate for synonymous substitution rate available for *O. vulgaris*. We determined historical peaks in transposon activity for each TE order separately, by grouping all JC distances into bins of size 0.01 and quantifying the cumulative sum of base pairs for all TEs of each order in each bin size. The bins that contained the highest number of base pairs for each TE order indicated periods of expansion.

### TE landscape

We identified hotspots and coldspots for all TEs in the *O. vulgaris* genome following Baril and Hayward (2022). Briefly, we divided each scaffold into 100kb windows and calculated the TE density within each window. This produced a frequency distribution of observations (Supplemental Material Figure S1) which we fit using a polynomial model to generate a smooth curve using the lm function in R (R Core Team 2024). We calculated the area under the curve (AUC) using the bayestestR package (Makowski et al. 2019) and from this we determined the 1% and 99% AUC cutoff values for coldspots and hotspots, respectively. We generated karyoplots using the KaryoplotR package (Gel and Serra 2017) to visualize the position of hotspots and coldspots along each scaffold. We also calculated the density of recently active TEs in each of the 100kb windows and visualized the position of recently active elements in the genome using karyoplots.

## Supplementary Material

Appendix I – Supplemental Methods and Appendix II – Supplemental Tables and Figures available online.

## Acknowledgements

We wish to thank Toby Baril for support with Earl Grey and the determination of TE cold and hotspots.

## Author Contributions

The authors confirm contribution to the paper as follows: *study conception and design*: SL, KC, TAE, MB; *data collection*: TAE, MAMA; *analysis and interpretation of results*: TAE, MB; *writing of the draft manuscript*: TAE, MB; *major comments and edits provided for the draft manuscrip*t: KC, SL. All authors reviewed the results and approved the final version of the manuscript.

## Data Availability

Data and code will be made available at the time of publication.

