## Supplemental Material for "Biological implications of a detailed repeat annotation in *Octopus vulgaris*"

**Appendix I - Supplemental Methods**

Reference TEs

References TEs for our detection and annotation process were collected from the following sources: a curated bivalve TE library (Martelossi et al., 2023), mollusc Penelope-like elements (Craig et al., 2021), TLEWI Tc1-Mariner elements (Puzakov et al., 2020), mollusc RTE LINE elements (Galbraith et al., 2022), bivalve TRIM elements (Satovic et al., 2019), *Donax trunculus* MITEs (Satovic and Plohl, 2013), Steamer Ty3-like elements (Arriagada et al., 2014; Metzger et al., 2018), *Haliotis discus hannai* Ty3-like elements (Lee et al., 2018), mollusc SINEs (Matetovici et al., 2016), Nin SINEs (Piskurek and Jackson, 2011), RUDI SINEs (Luchetti et al., 2016) and others (Pierce et al., 2016; Satovic and Plohl, 2017; Wang et al., 2008).

**Appendix II - Supplemental Tables and Figures**

Supplementary Table 1. Detailed breakdown of repetitive DNA in the xcOctVulg1.2 *Octopus vulgari***s** genome.

| **Repeat Type** | **Subdivision** | **Coverage (bp)** | **Coverage (%)** | **Consensus Sequences/Families** |
| --- | --- | --- | --- | --- |
| Transposable Element Total |  |  | 40.19% | 3343 |
| Retrotransposon |  |  | 32.36% | 2286 |
|  | LINE | 906379368 | 12.97% | 633 |
|  | RTE | 363367447 |  | 176 |
|  | Dong-R4 | 106052287 |  | 51 |
|  | CR1 | 92734607 |  | 169 |
|  | L1-Tx1 | 62185422 |  | 80 |
|  | Unclassified LINE | 56276355 |  | 11 |
|  | L1 | 22957682 |  | 60 |
|  | Hero | 10836547 |  | 14 |
|  | CRE | 8222957 |  | 8 |
|  | Proto2 | 1413969 |  | 13 |
|  | I | 539601 |  | 26 |
|  | R2 | 111411 |  | 1 |
|  | RTE-X | 89048 |  | 24 |
|  | SINE | 328452608 | 11.72% | 1317 |
|  | Unclassified SINE | 249689970 |  | 1279 |
|  | tRNA-Deu | 60937931 |  | 17 |
|  | 5S | 9831953 |  | 4 |
|  | tRNA-CORE | 4357417 |  | 2 |
|  | tRNA-CORE-RTE | 3136658 |  | 4 |
|  | U | 369490 |  | 1 |
|  | U-L1 | 52753 |  | 1 |
|  | Other SINE | 76436 |  | 9 |
|  | Penelope-like | 191478272 | 6.83% | 190 |
|  | Penelope | 183798989 |  | 172 |
|  | Other-Penelope-like | 7679283 |  | 18 |
|  | LTR | 21557928 | 0.77% | 141 |
|  | Ty3-like | 14677594 |  | 116 |
|  | Otner LTR | 6880334 |  | 25 |
|  | Other Retrotransposon | 1523113 |  | 5 |
| DNA Transposon |  | 219212292 | 7.82% | 1054 |
|  | TIR | 190602669 | 6.8% | 963 |
|  | Tc1-Mariner | 119138219 |  | 362 |
|  | hAT | 16560276 |  | 134 |
|  | PiggyBac | 15558031 |  | 36 |
|  | Mutator | 13868679 |  | 55 |
|  | Unclassified TIR | 12404772 |  | 123 |
|  | CACTA-Mirage-Chapaev | 4136930 |  | 22 |
|  | Merlin | 2185638 |  | 18 |
|  | Sola1 | 2041902 |  | 7 |
|  | Kolobok | 1903292 |  | 22 |
|  | PIF-Harbinger-ISL2EU | 941386 |  | 46 |
|  | Ginger1 | 824851 |  | 1 |
|  | Ginger2 | 526085 |  | 1 |
|  | Academ | 200619 |  | 48 |
|  | MITE | 186477 |  | 74 |
|  | Sola2 | 98818 |  | 7 |
|  | Other TIR | 26694 |  | 7 |
|  | Helitron | 28598833 | 1.02% | 84 |
|  | Other DNA | 10790 |  | 7 |
|  | Unclassified Transposon | 45628 |  | 3 |
| Simple |  | 208872170 | 7.45% |  |
| Low complexity |  | 9843447 | 0.35% |  |
| Satellite |  | 87464451 | 3.12% | 80 |
| Unknown |  | 423466180 | 15.12% | 1102 |
| **Repeat Total** |  | **1855283536** | **66.25%** | **4525** |

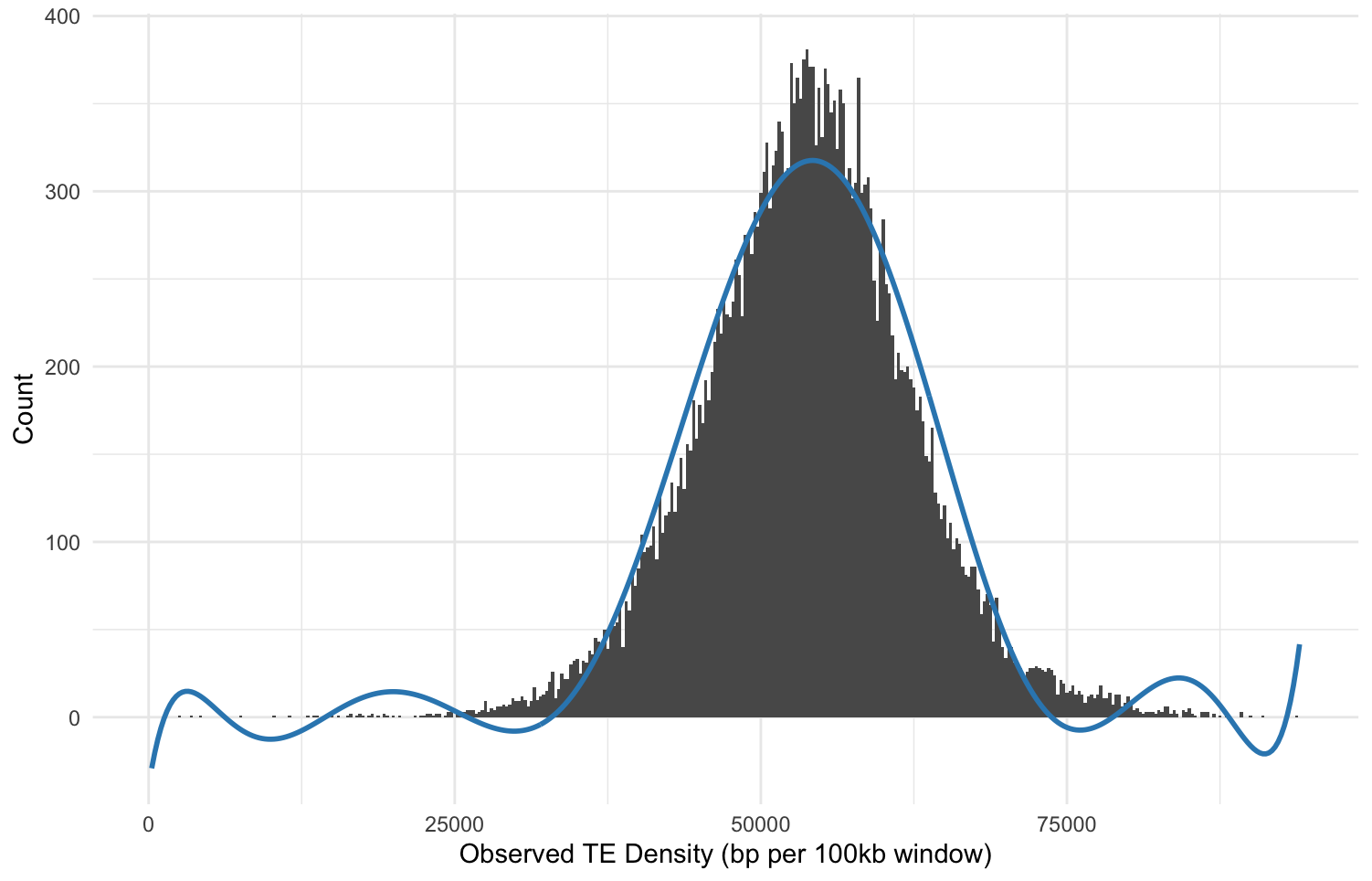

**Figure S1.** Frequency distribution of 100kb windows containing TEs in *O. vulgaris* genome. The blue line is the polynomial model used to generate a smooth curve for calculation of TE hot and coldspots (See Materials and Methods).
